# The polygenic strategies of *Botrytis cinerea* for virulence across eight eudicot host species

**DOI:** 10.1101/2024.08.19.608699

**Authors:** Céline Caseys, Daniel J. Kliebenstein

## Abstract

Diverse qualitative and quantitative genetic architectures can successfully influence fungal virulence and host range. To model the quantitative genetic architecture of a generalist pathogen with an extensive host range, we conducted a genome-wide association study (GWAS) of the virulence of *Botrytis cinerea* across eight hosts. This approach revealed 4772 candidate genes, about one-third of the *Botrytis* genome, contributing to virulence with small effect sizes. The candidate genes are evenly spread across the core chromosomes with no indication of bipartite genomic architecture. The GWAS-identified polymorphisms and genes show that *B. cinerea* relies on genetic variants across hundreds of genes for infecting diverse hosts, with most genes influencing relatively few hosts. When pathogen genes associated to multiple hosts, they typically influenced more unrelated than related host species. Comparative genomics further suggested that the GWAS-identified genes are largely syntenic with other specialist Botrytis species and not novel to *B. cinerea*. Overall, *B. cinerea*’s generalist behavior is derived from the sum of the genome-wide genetic variation acting within gene networks that differentially coordinate the interaction with diverse hosts.

## Introduction

Plant pathogens are major threats to global food security as they cause significant yield losses (Strange & Scott, 2005; Singh, BK *et al*., 2023). However, most plant-microbe interactions do not result in disease, either due to the success of the plant defenses or the microbes’ lack of virulence. Those defense/virulence molecular strategies are encoded in the host and microbes genomes and the genetic variation in plants interacts with the genetic diversity within pathogens (Laine *et al*., 2011; Barrett *et al*., 2015). It is critical to understand how the genetic variation in both organisms leads to disease and the pathogen’s ability to cause disease in single to multiple hosts, also known as host range (Barrett *et al*., 2009; Morris & Moury, 2019).

For some plant-pathogen interactions, the genetics determining the interaction relies on a few genetic variants of large effect creating qualitative variation in the virulence outcome (Dong *et al*., 2015; Möller & Stukenbrock, 2017). These large-effect genes can be present on accessory chromosomes such as in *Alternaria alternata*, close to repetitive elements such as in *Magnaporthe oryzae* or close to recombination hotspots such as in *Melampsora lini* (Li et al., 2020). Often clusters of these genes are associated together and evolve more rapidly than the rest of the genome leading to the appearance of a two-speed genome (Dong *et al*., 2015; Torres *et al*., 2020). In other species, such as *Blumeria graminis*, host specialization genes don’t appear to cluster but are boosted by a high duplication rate and a dynamic secretome (Frantzeskakis *et al*., 2018). Horizontal transfer of those effector genes or accessory chromosomes between pathogen species can further influence the genetics determining host range (Mehrabi *et al*., 2011).

Beyond large effect genes, the genetic architecture of virulence and host range can also be quantitative involving broad sets of genes (Poland *et al*., 2009), each contributing to the outcome with small effect sizes (Liao *et al*., 2022). The broad sets of genes influencing virulence and host range in the quantitative systems tend to involve an equally diverse set of molecular mechanisms. For example, in *Zymoseptoria tritici*, quantitative pathogenicity genes are hotspots for diversifying selection (Amezrou *et al*., 2024), with heterochromatin accessibility and transposable elements also playing an important role in creating variation (Fraser & Whitehall, 2022). In *Sclerotinia sclerotiorum*, quantitative effects on virulence and host range involve the production of distinct small RNAs (Derbyshire *et al*., 2019), transcriptional reprogramming (Allan *et al*., 2019; Kusch *et al*., 2021), and alternative splicing (Ibrahim *et al*., 2021). How the quantitative architectures vary with the lifestyle (biotroph vs necrotroph) and host range of plant pathogens remains poorly understood. We used *Botrytis cinerea* to investigate how quantitative genetics influence virulence and host range in a generalist necrotroph.

*Botrytis cinerea* (Grey mold, Botrytis thereafter) is a plant pathogen that causes damage to over a thousand plant species from more than 600 genera across a wide phylogenetic spectrum, ranging from mosses to monocots and eudicots, including major crops (Elad *et al*., 2016; Singh, R *et al*., 2023). Botrytis threatens food security both pre- and post-harvest, with the ability to attack diverse hosts and organs within each host such as the leaf, flower, fruits, and stem (Dean *et al*., 2012). The mechanism(s) enabling *Botrytis cinerea* to be such a successful generalist are not clear given that other Botrytis species tend to be specialists (Navaud *et al*., 2018; Valero-Jiménez *et al*., 2019; Valero-Jiménez *et al*., 2020; Garfinkel, 2021). Thus, this system has the potential to better understand how generalist plant pathogens may evolve and the mechanisms involved.

As a necrotroph, Botrytis kills and feeds on cells it attacks (Veloso & van Kan, 2018; Bi *et al*., 2023) using a variety of polymorphic mechanisms. The genetic architecture of Botrytis virulence is highly quantitative with over a hundred genes having a validated role in modulating virulence and potentially host range (Williamson *et al*., 2007; Nakajima & Akutsu, 2013; Mbengue *et al*., 2016; Veloso & van Kan, 2018; Cheung *et al*., 2020; Bi *et al*., 2023; Singh, R *et al*., 2023). These genes include signaling genes in molecular pathways such as cAMP (Kronstad, 1997), G proteins (Li *et al*., 2007), calcium-dependent (Lange & Peiter, 2020), and MAPK cascades (Lange & Peiter, 2020) that are historically associated to virulence in fungi with other lifestyles (e.g. biotrophs). Part of the interaction with the host is also mediated through small RNAs that interfere with the host’s resistance mechanisms (Weiberg & Jin, 2015). Other major parts of the Botrytis virulence toolbox are toxins (such as botrydial) (da Silva Ripardo-Filho *et al*., 2023), cell wall degrading enzymes, and cell death-inducing proteins (Mbengue *et al*., 2016; Bi *et al*., 2023). Further contributing to the host cell death are genes modulating redox, pH, and the active generation of reactive oxygen species (ROS) (Heller & Tudzynski, 2011; Newman & Derbyshire, 2020). In addition to the above offensive mechanisms, Botrytis also has numerous and diverse defensive mechanisms to resist the various phytochemical defenses produced by the broad range of host plants (Stefanato *et al*., 2009; Pedras *et al*., 2011; Westrick *et al*., 2021; Kuroyanagi *et al*., 2022; Bulasag *et al*., 2023). How this diversity of molecular mechanisms contributes to influencing virulence across the broad-host range of *Botrytis cinerea* is presently unclear.

With over a thousand plant hosts (Elad *et al*., 2016; Singh, R *et al*., 2023), it is imperative to begin modeling how many Botrytis genes contribute to the virulence on a single host versus multiple hosts. Is this host range due to genes specific to hosts or genes functioning across diverse hosts using host specific alleles? For example, in Polygalacturonase 1 (PG1) different alleles might degrade various pectin structures across host families (Rowe et al. 2007). However, the relative contribution of host specific genes or host specific alleles remains to be catalogued. Furthermore, identifying the genes influencing Botrytis virulence across diverse host plants is crucial to understanding the origins of its extreme polyphagy within the Botrytis genus (Navaud *et al*., 2018; Mercier *et al*., 2019). Such polyphagy could result of the evolution of novel genes or retention/collection of host-specific genes found in other Botrytis species. Additionally, mapping the genetic architecture of virulence and host range will aid in detecting signs of clustering within the genome that could contribute to the broad host range.

To begin querying how the host range and virulence are determined in Botrytis, we used previous measurements of disease caused by a population of 96 diverse Botrytis isolates across an array of eudicots including tomato, sunflower, lettuce, chicory, endive, turnip, Arabidopsis, and soybean (Caseys *et al*., 2021). In this work, we use these phenotypic measurements to identify the Botrytis genes that may shape host susceptibility. Combining the phenotypic measurements with the genomic variation in the Botrytis population (Atwell *et al*., 2015; Atwell *et al*., 2018; Soltis *et al*., 2019; Zhang *et al*., 2019; Krishnan *et al*., 2023), we mapped and analyzed the genetic architecture of host preferences across Botrytis isolates using genome-wide association study (GWAS).

## Materials and methods

### Quantifying plant-pathogen interactions

A collection of 96 isolates of *B. cinerea* was used to infect 85 plant genotypes from eight different eudicot species. The pathogen isolates originated largely from California with some isolates coming from elsewhere around the world (Caseys *et al*., 2021). The plant genotypes (Table S1) cover eight species, including *Arabidopsis thaliana,* and seven crops. For each crop, there are six genotypes representing accessions either from wild or landrace origin and six genotypes are inbred lines or cultivars (Caseys *et al*., 2021). The crop species were selected both within the Rosids and the Asterids. For the Rosids, *Brassica rapa* (turnip) and *A. thaliana* are in the Brassicales while *Glycine max* (soybean) is in the Fabales. Within the Asterids a range of phylogenetic distances are present. *Solanum lycopersicum* (domesticated tomato) and *Solanum pimpinellifolium* (wild tomato) are in the Solanales. Within the Asterales, two sister chicory species *Cichorium intybus,* and *Cichorium endivia* were compared to *Lactuca sativa* (domesticated lettuce), *Lactuca serriola* (wild lettuce), and *Helianthus annuus* (sunflower).

Detached leaf assays were performed with six-fold replication across two independent experiments by infecting adult leaves with 4ul drops of 50% grape juice containing 40 Botrytis spores (Caseys *et al*., 2021). The lesion areas were quantified 72 hours post-infection by taking pictures of lesions, counts of pixels within the lesion, and conversion into square centimeters. To account for the effects of experimental factors, the lesion areas were analyzed with linear mixed models and for each genotype by isolate, model-corrected least-square means were calculated (Caseys *et al*., 2021).

### Genome-wide association studies (GWAS)

To map the genetic variation in Botrytis influencing the lesion area on the 85 plant genotypes infected with the 96 Botrytis isolates, a GWAS approach was applied. For the genotyping information, we used a previously published set of 271,749 single nucleotide polymorphisms (SNPs) at a minor allele frequency of (MAF) 0.2 with less than 20% missing data mapped (Atwell *et al*., 2015; Atwell *et al*.; Soltis *et al*., 2019) to the B05.10 genome ASM83294v1 assembly (Van Kan *et al*., 2017).

No evidence of host specialization or population structure was detected among the 96 isolates of this haploid fungus (Atwell *et al*., 2018; Caseys *et al*., 2021). Given the quantitative nature of Botrytis virulence (Soltis *et al*., 2019; Caseys *et al*., 2021), a Bayesian sparse linear mixed model (BSLMM) using Markov chain Monte Carlo algorithm implemented in GEMMA was run for lesion area on each plant genotype (Zhou & Stephens, 2012). BSLMM was chosen because it concurrently models two effect size distributions: a polygenic architecture made of loci of small effect sizes and an oligogenic architecture allowing for some loci of modest to larger effect sizes (Zhou *et al*., 2013). A standardized relatedness matrix was included in the model to account for genetic similarity and any population structure. For each trait, 20 independent runs with 500000 burn-in and 5000000 iterations with recording every 10 steps were performed. For these models, we constrained the parameter h, an estimate of the narrow-sense heritability, between 0.1 < h < 0.9. All studies to date (Zhang *et al*., 2017; Soltis *et al*., 2019; Caseys *et al*., 2021) have shown heritability rates within this range for Botrytis virulence.

BSLMM outputs both SNP estimates of effect size and significance, estimated as the posterior inclusion probability (PIP). For each SNP, the median of the effect size and PIP distribution of the 20 runs was used for subsequent analyses. To estimate the null distribution of the PIP, 10 random permutations were performed for each host species revealing a common trend across species (Figure S1).

From the random permutation, it was estimated that PIP values larger than 1.7 x10^-4^ are equivalent to a 5% chance of false positives. The BSLMM results were then filtered using this 5% chance of false positive threshold and further analyzed for multivariate and local false sign rate (LFSR).

To incorporate the phenotypic information potentially provided by lesion area measured on up to 12 plant genotypes per host species, we implemented a multivariate approach to the BSLMM using multivariate adaptive shrinkage (MASH) (Urbut *et al*., 2018). MASH analyzes the combined effects and standard errors of GWAS-associated SNPs across the phenotypes while accounting for arbitrary correlations among conditions. This multivariate approach did not result in significant results when combining the GWAS across all the host species indicating the absence of universally significant SNPs. MASH models were then applied to each host using all the independent genotypes per host and covariance matrices and LFSR posterior summaries were extracted. Those LFSR are analogous to the false discovery rate. SNPs with LFSR <0.05 were considered as significant for the given host genotype and significances consolidated within each species.

To test for relationships in the SNP significances across the host genotypes, we conducted a hierarchical clustering across the 85 genotypes using the “complete” agglomeration method. This method calculates the maximum Euclidean distance between clusters before merging. The significance of the branches was assessed with the r package pvclust over 20000 bootstraps. Branches were declared significant if the approximately unbiased p-values (AU) estimated using bootstraps were larger than 95%.

To obtain a genome-wide view of the distribution of the significant SNPs and their local effect, an effect size sliding window analysis was performed. Given the level of linkage disequilibrium (Atwell *et al*., 2018; Mercier *et al*., 2021) within Botrytis and the potential for multiple significant SNPs within genes, we used a sliding window of 1kb, with steps of 500bp. Within each window, we summed the local effect size estimates for each SNP within the window.

### Candidate gene analyses

Given the average haplotype diversity and linkage disequilibrium along the Botrytis genome giving approximately gene-level resolution for mapping (Atwell *et al*., 2018; Mercier *et al*., 2021), we consolidated the significant SNPs identified by grouping them into their associated genes for further analyses. Due to the gene density in the Botrytis genome and to take into account regulatory regions, SNPs within 500bp of the start or end of the gene were consolidated as being part of the gene. The count of significant SNPs per gene and their location can be found in Table S2.

To reveal how candidate Botrytis genes were shared across host plant species, a Venn diagram with seven species (excluding *A. thaliana* which has only one genotype) was drawn with the R library “Venn”. This representation with 7 species is the most complete option available. Furthermore, hierarchical clustering of the candidate genes across hosts was run as the SNP analysis described above.

To estimate the proportion of the candidate genes that might have recently evolved within *B. cinerea*, we performed comparative genomics analysis with 7 Botrytis species. *B. cinerea* B05.10 genes orthology to *B. fragariae* (Wu *et al*., 2021), *B. aclada*, *B. deweyae*, *B. porri*, *B. hyacinthi* and *B. sinoallii* (Valero-Jiménez *et al*., 2019; Valero-Jiménez *et al*., 2020) were called by OrthoMCL in KBase (Arkin *et al*., 2018). *B. fragariae* is found in strawberry fields in Germany and South East United States (Rupp et al., 2017)*. B. deweyae* is an endophytic facultative necrotroph, found on one plant species (polyphagy index=1). All other species are necrotrophs infecting monocots infecting 3-9 host species and polyphagy index between 1.4 and 3.2 (Mercier *et al*., 2019).

To estimate putative mechanistic connections among candidate genes, gene co-expression networks were drawn using 16 hours post inoculation transcriptomes measured from these same Botrytis isolates infected on *Arabidopsis thaliana* Col-0 leaves (Zhang *et al*., 2017; Zhang *et al*., 2019). In the networks, vertices connect transcripts with a Spearman’s rank correlation of ρ>0.75 and p-value<0.05. Networks were constructed with the R package “network” and plotted with the gplot function of the package “sna” with vertices placed according to the Fruchterman-Reingold algorithm.

Genes were annotated with gene names, genomic location, protein features, and function prediction as present on fungidb.org (Besenko et al. 2018) in January 2024. ncRNAs were annotated with tRNAscan-SE (Chan & Lowe, 2019). Further annotations were added manually for genes part of the surfactome (Escobar-Nino et al 2021) and secretome (González-Fernández *et al*., 2015).

## Results

### Polygenic architecture of virulence across host species

A detached leaf assay measuring the disease outcome of a population of Botrytis isolates on diverse eudicot hosts was conducted in a previous study (Caseys *et al*., 2021). The measurement of the lesion area associated with Botrytis necrotrophic growth on leaves is genetically heritable and variable across the 85 plant genotypes tested across eight plant species (Figure S2A). The percentage of variance in the lesion area explained by the genetic variation between Botrytis isolates ranged from 21% (*A. thaliana*) to 51% (*C. endivia*) (Figure S2A). We used these measures of lesion area across 96 Botrytis isolates (Figure S2B) to conduct a GWAS and map virulence loci and architecture across the diverse host plants.

Given previous work on Arabidopsis (Zhang *et al*., 2019; Krishnan *et al*., 2023), tomato (Soltis *et al*., 2019), and genome scans (Mercier *et al*., 2021), we anticipated predominantly small effect virulence loci with a chance for occasional loci with moderate effect sizes. To match the GWAS model to this architecture, we used BSLMM, which can estimate genome-wide small effects (a polygenic architecture modeled by linear mixed model) and additional moderate effects (an oligogenic architecture modeled by Bayesian regression model). Using the resulting data, we first quantified the genetic architecture of pathogen virulence across all the host genotypes with four BSLMM hyper-parameters: the proportion of heritability explained by the Bayesian regression model (rho), the percentage of variance explained by both models (PVE), the number of variants with larger effect (gamma) and the proportion of PVE explained by variants with larger effects (PGE).

This revealed that the genetic architecture of the lesion area across all plant genotypes was quantitative and complex. Using BSLMM, it is possible to estimate a rho parameter that can vary from polygenic (rho=0) to oligogenic (rho=1) and provide an empirical assessment of the genetic architecture (Zhou *et al*., 2013). Across all 85 different plant hosts, there was an average of rho=0.5 with a range of rho=0.41 (*C. intybus* PI652041, Figure 1A) to rho=0.58 (*C. intybus* PI651945, Figure 1A). This suggests the genetic architecture is largely similar across all hosts and either slightly oligogenic or polygenic, with no strong model directionality and variation across genotypes of a host (Figure 1A). The BSLMM also estimated the percentage of phenotypic variance explained by the possible genetic architectures. Considering the effect of all 271481 SNPs in the genome explained 16% (*L. sativa* LJ10335) to 53% (*C. intybus* PI652021) of the phenotypic variance (Figure 1B) across the 96 Botrytis isolates. This range of percentage of phenotypic variance is likely due to variation in the influence of plant genetics across the species (Figure S2A). When modeling for moderate effects, the oligogenic model consistently found 10 to 15 SNPs (Figure 1C) influencing virulence on each genotype.

**Figure 1:**
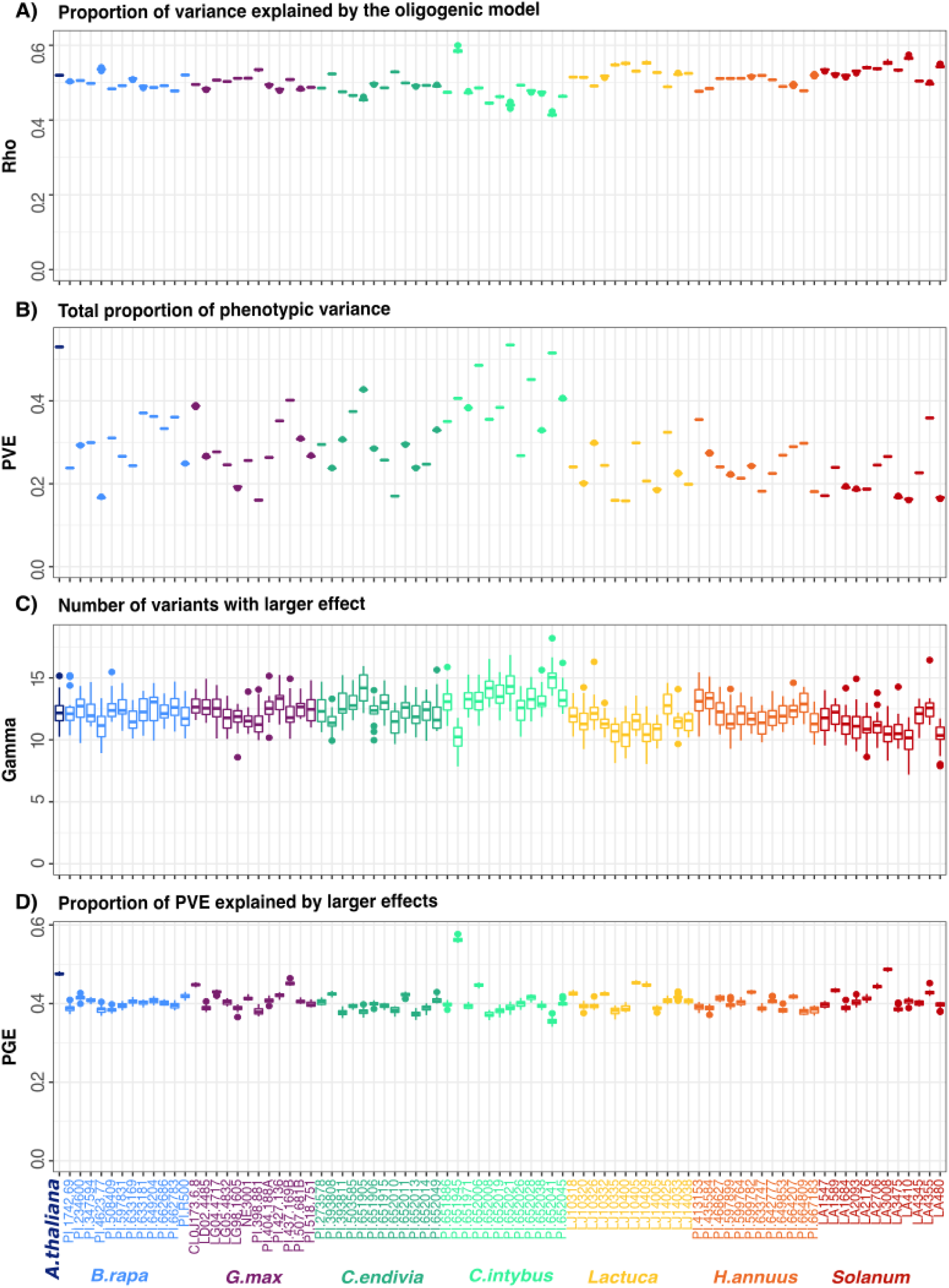
Genetic architecture of Botrytis quantitative virulence on 85 host plants. For each BSLMM hyperparameter, the mean of the recorded iterations for the 20 separate BSLMM runs is provided. The plant accessions on which the lesion area was measured (phenotypes) are colored by species. A) Rho describes the proportion of genome-wide small effects (rho=0 polygenic) to relatively larger effects (rho=1 oligogenic). B) Total percentage of phenotypic variance explained (PVE) by all SNPs included in the model. C) Number of SNPs with relatively larger effects estimated by the oligogenic model. D) Percentage of the PVE explained by the SNPs with relatively larger effects.

Inclusive of lesion measurements for all 85 hosts, 27481 SNPs were significantly (LFSR<0.05) associated to the lesion area. We first tested if genomic features (3’UTR, 5’UTR, CDS, Intergenic, Intron) were enriched in significant SNPs (Figure S3). To create a random null distribution, we selected 27481 random SNPs that match the MAF of the significant SNPs and quantified the associated genomic features. Repeating this one hundred times created permutation sets of genomic features. This showed that the significant SNPs represented a null distribution and had no significant enrichment in genomic features. To test if the significant SNPs were evenly distributed across the genome or potentially showed any local clustering of virulence loci (Torres *et al*., 2020), we mapped the position and effect of the significant SNPs across the Botrytis genome using 1kb windows which matches the average distance of linkage disequilibrium decay (Atwell *et al*., 2018). This revealed an even distribution of significant loci across the entire genome (Figures 2A, S4) having mainly small effect sizes (Figure S5). The local effect size displays a long-tail distribution within each host species, with a few regions having a larger albeit still small effect than the rest of the genome (Figure S5). There was no overlap across host species in these regions. Further, when comparing these outliers and most significant SNPs across species, there are relatively few common SNPs across hosts, with most significantly associated SNPs being largely unique to a host or shared with one other host (Figure 2B). Only 94 SNPs are associated with Botrytis virulence across 5 or more species (Figure 2B) and are spread across the genome (Figure S4).

**Figure 2:**
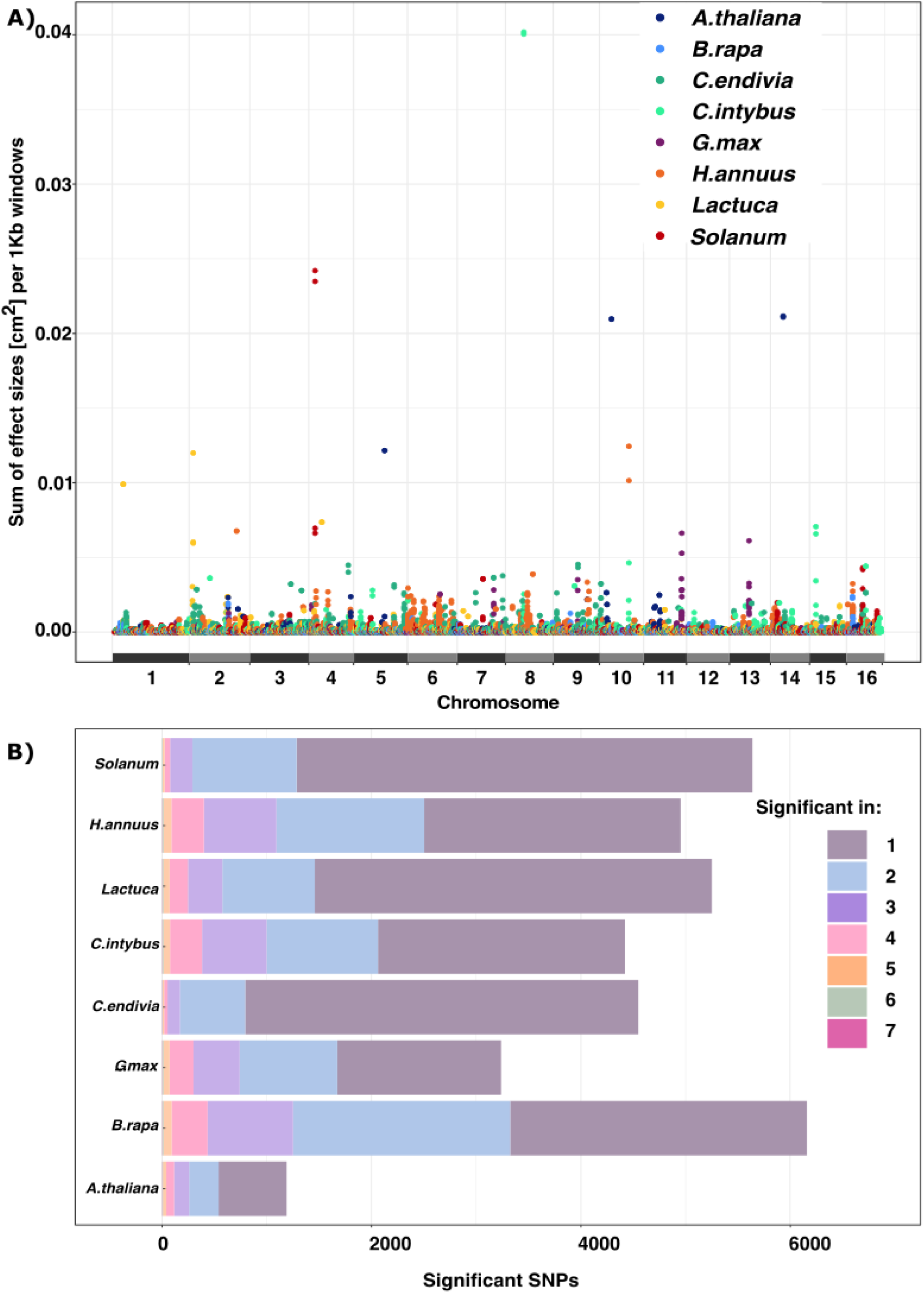
Virulence-associated SNPs are spread across the Botrytis core chromosomes and largely associated to a single host. A) Manhattan plot of the effect size of SNPs significantly associated (LFSR<0.05) to lesion area [cm^2^]. The effect sizes are summed within 1kb window. The colors indicate the host with the maximal effect size for each window. B) Counts of SNPs significantly associated (LFSR<0.05) to lesion area for each host species. The colors indicate whether the SNP is uniquely associated to a host (1) or shared with up to 7 hosts (7). No gene was identified in all eight hosts.

At the phenotypic level, a previous analysis suggested similarities in the virulence across genotypes within a host but little phylogenetic relationship between host species (Caseys *et al*., 2021). To test how the significant SNPs may associate across host genotypes and assess if there was a signal from the phylogenetic relationships across the host species, we conducted a hierarchical clustering of all SNPs based on their effect within a host genotype (Figure S6). This revealed that the Botrytis causal SNPs find a common signal across genotypes within a host species. To utilize this common signal, we merged the significant SNPs across the host genotypes within a species to give a single image of the virulence architecture of Botrytis per species.

### A large proportion of the Botrytis genome is involved in virulence

To transition from individual SNPs and identify mechanistic signals, we used the SNP’s positions in the B05.10 ASM83294V1 reference genome (Van Kan *et al*., 2017) to map significant SNPs to candidate genes. This revealed a total of 4772 genes and 62 transfer, ribosomal and spliceosomal RNAs (Table S2). The candidate genes contained between 1 and 90 significant SNPs (Table S2) with an average of 3 (median of 2) significant SNPs per gene. With 14236 coding genes in the reference genome (Van Kan *et al*., 2017), the candidate genes associated with virulence effects in one or more host species account for 33% of the coding genes. Out of the 4772 genes associated with significant SNPs, 3670 genes have functional annotations (Figure 3, Table S2). Given the large number of candidate genes and the sparsity of specific annotations in the Botrytis genome, no enrichment of gene ontology (GO) or protein domains was detected.

**Figure 3:**
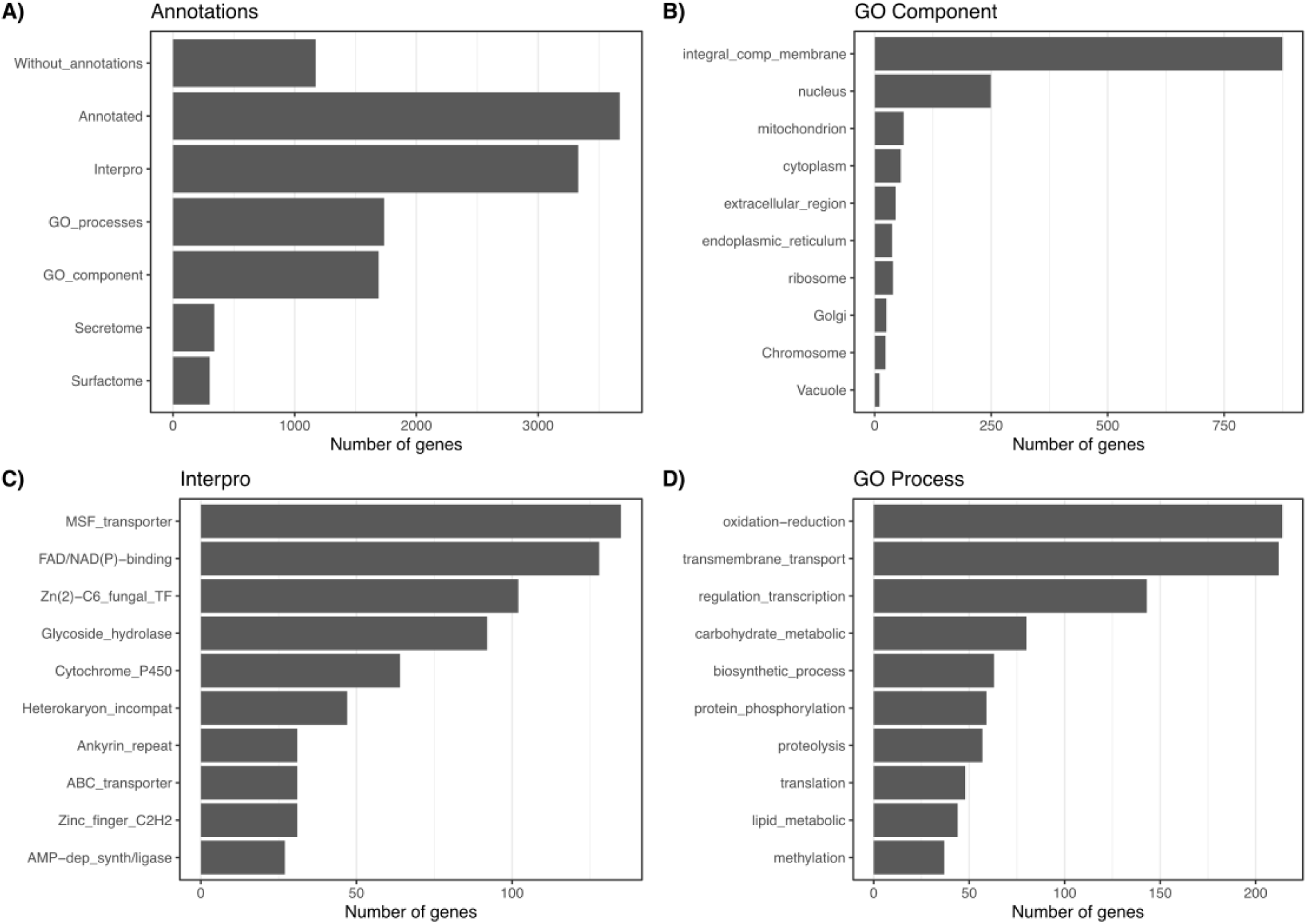
The Botrytis genes associated to virulence cover a large range of functions. This figure provides an overview of the annotations of candidate genes presented in Table S2. A) Number of candidate genes with functional annotations. B) Top 10 gene ontologies for cellular components. C) Top 10 Interpro protein domain annotations D) Top 10 gene ontologies for biological processes.

Instead, the candidate genes follow gene functional diversity across the whole genome. Among functional categories known for their role in virulence, the interaction with the host, cellular stress responses, regulation of the transcription, and biosynthetic processes are within the top 10 functional annotations (Figure 3). Proteins that are integral components of the membrane are the top cellular component category (Figure 3B, Table S2). The potential key role of transmembrane transport (Figure 3D) is driven by numerous MSF and ABC transporters (Figure 3C). The role of cellular stress response is highlighted by oxidation-reduction (Figure 3D) and FAD/NAD(P) binding annotations (Figure 3C). The potential key role of regulation of the transcription (Figure 3D) is driven by the numerous Zn(2)-C6 and C2H2 transcription factors (Figure 3C).The role of biosynthetic processes (Figure 3D) is also associated with the numerous glycoside hydrolase and cytochrome P450 (Figure 3C). Using previous mechanistic studies, 72 genes identified by the GWAS have been validated for their role in Botrytis growth, development, or virulence (Table S2).

One possible model that could explain *Botrytis cinerea*’s generalism is that it could contain a collection of novel genes evolved to provide virulence across numerous hosts. To test if the candidate genes influencing quantitative virulence are unique to *Botrytis cinerea* or are present within other narrower host-range Botrytis species (Valero-Jiménez *et al*., 2019; Valero-Jiménez *et al*., 2020), we performed comparative genomics, calling orthology groups across seven Botrytis species from clade 1 and 2 (Garfinkel, 2021). This analysis revealed that 3727 genes are fully shared across seven Botrytis (Figure 4) species while 794 genes are partially syntenic, being shared with one to five other Botrytis species. Only 210 genes have an indication that they might be specific to *B. cinerea* B05.10 isolates. To create a null distribution of orthologs, we randomly selected a thousand sets of the *B. cinerea* orthologs (Figure S7A). This showed that the genes identified by GWAS are not enriched for genes specific to *B.cinerea* (Figure S7B). When comparing the observed (identified by GWAS) and expected distribution of genes shared with the other Botrytis species, candidate genes are less likely than by chance to be shared with *B. aclada*, *B. hyacinthi,* and *B. porri* (Figure S7C).

**Figure 4:**
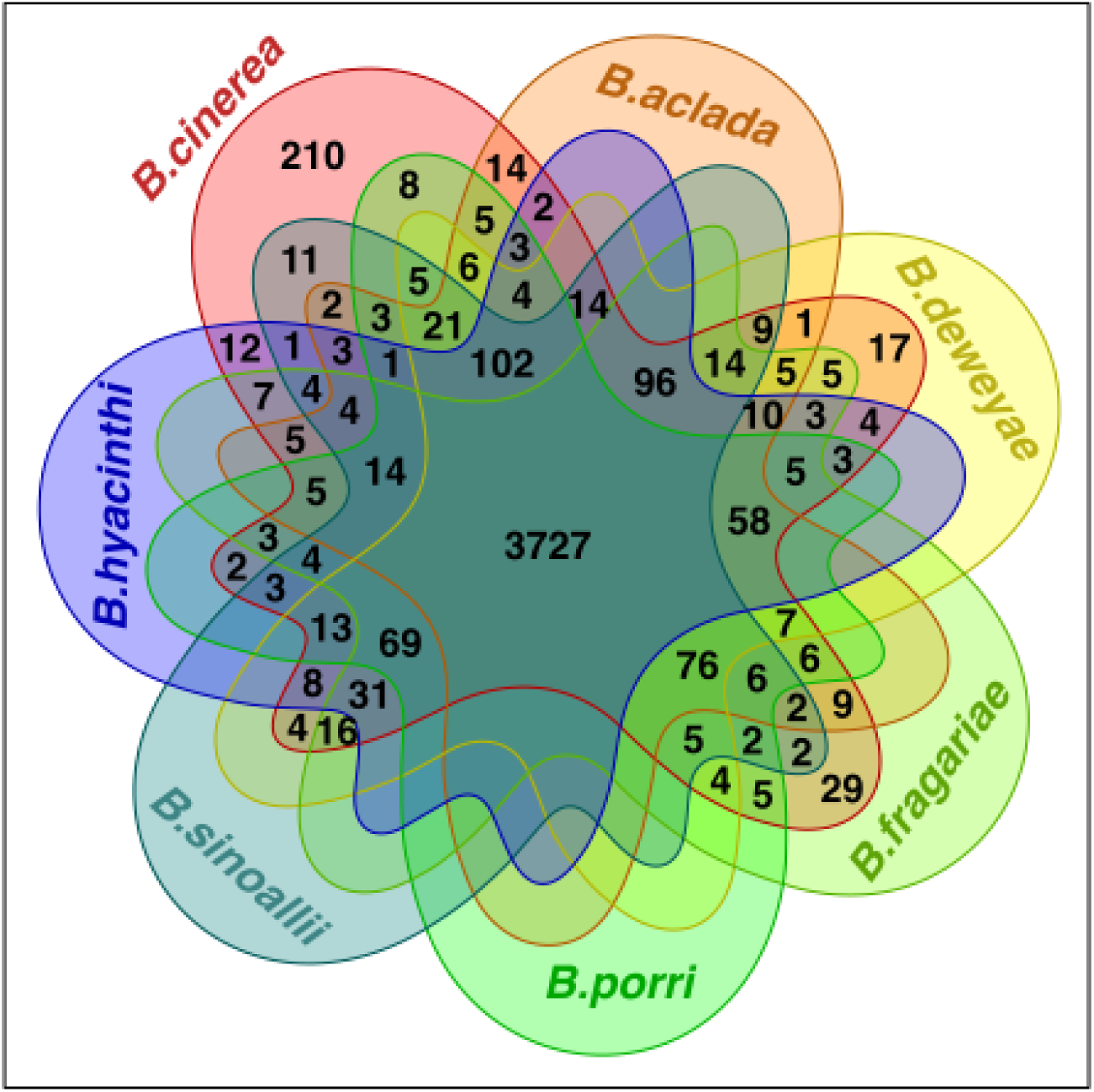
The *B.cinerea* genes associated to virulence are highly syntenic with other Botrytis species. Venn diagram with the count of *B. cinerea* B05.10 (clade 1, polyphagy index=54.1) genes shared with *B. fragariae* (clade 1, polyphagy index=1), *B. aclada* (clade 2, polyphagy index=1.4), *B. deweyae* (clade 2, polyphagy index=1), *B. porri* (clade 2, polyphagy index=1.4), *B. hyacinthi* (clade 2, polyphagy index=3.2) and *B. sinoallii* (clade 2, polyphagy index=1.4).

#### The genetics of Botrytis host range

Given the high proportion of the genome potentially involved in virulence, two polygenic models could explain virulence across hosts. First, the virulence across hosts could involve a limited set of genes with a large number of alleles that provide different host specificities. Under this model, there would be a common set of virulence genes across hosts with host-specific SNPs. Alternatively, virulence could be explained by a diverse set of genes that influence host range with unique polymorphisms. In this model, SNPs and genes would be equally host-specific. Surveying the candidate genes across the hosts showed that, like the SNPs (Figure 2B), the genes are largely unique to one or two host species (Figure 5A). Few genes are associated with a majority of hosts (Figure 5), with ten genes shared across seven species (Figure 5A, Table S2), and 47 genes shared across six species (Figure 5A, Table S2). This suggests that Botrytis virulence across these hosts is influenced by a wide set of genes with unique SNPs rather than a common set of genes with host-specific SNPs.

**Figure 5:**
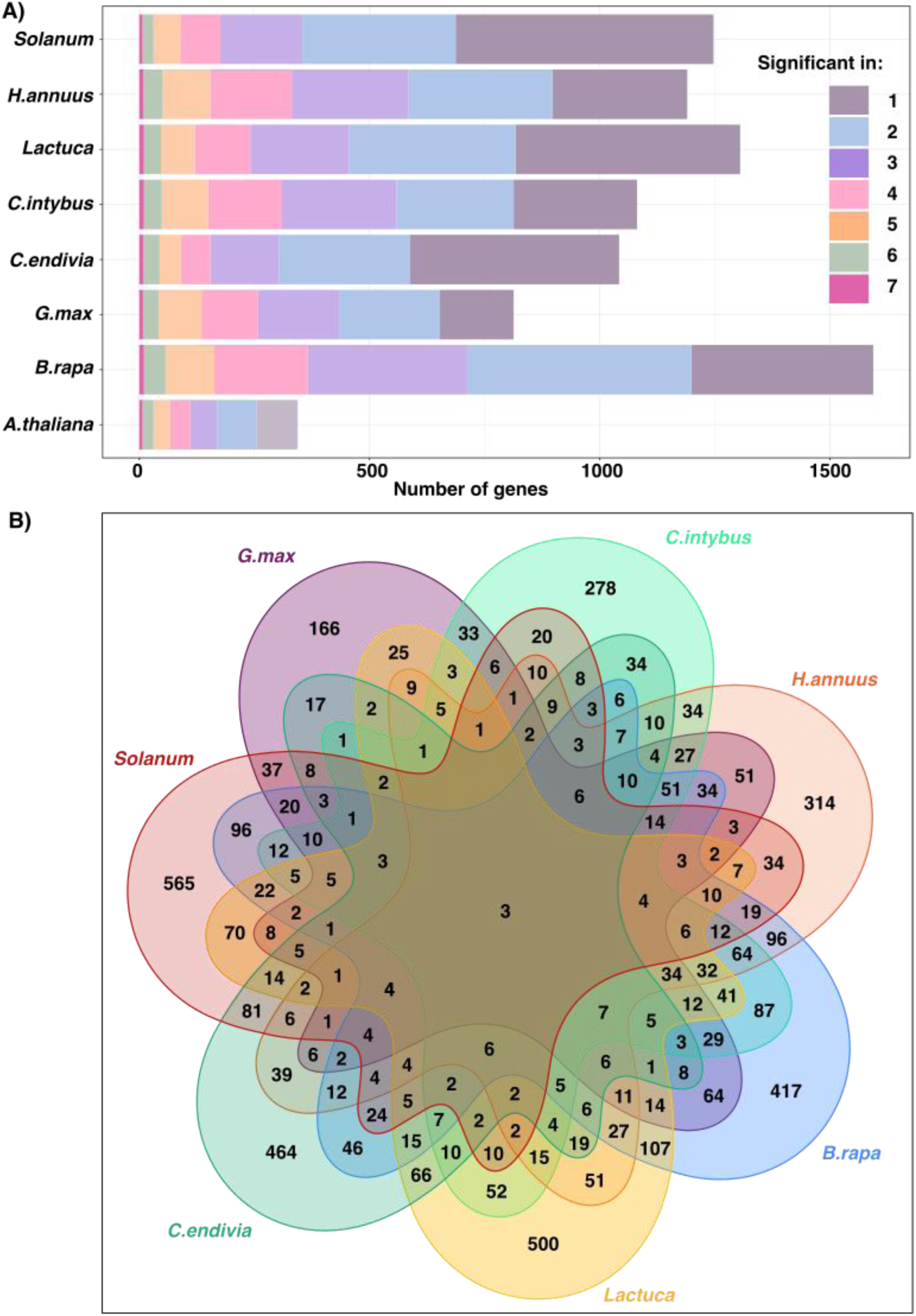
The Botrytis genes associated to virulence are largely shared across a few hosts. A) Number of candidate genes associated to the lesion area on eight eudicot hosts. The gene counts are colored based on association to 1 to 7 hosts. B) Venn diagram of candidate genes associated to lesion area on seven hosts. *A. thaliana* was excluded because only a single genotype (Col-0) is reported while for the other species 12 genotypes are reported.

To test how the candidate genes may influence virulence across the phylogenetic relationship of the hosts, we assigned the candidate genes to the phylogenic tree of the hosts. This representation of the evolutionary history of the hosts revealed that genus level (e.g. Cichorium Figure 6A) and order (e.g. Brassicales, Figure 6A) shared more genes than the clade level (e.g. Rosids and Asterids, Figure. 6A). However, given the complex patterns of shared genes across the hosts (Figure 5B), this analysis only accounts for 83 genes associated to multiple hosts that share a phylogenetic relationship.

**Figure 6:**
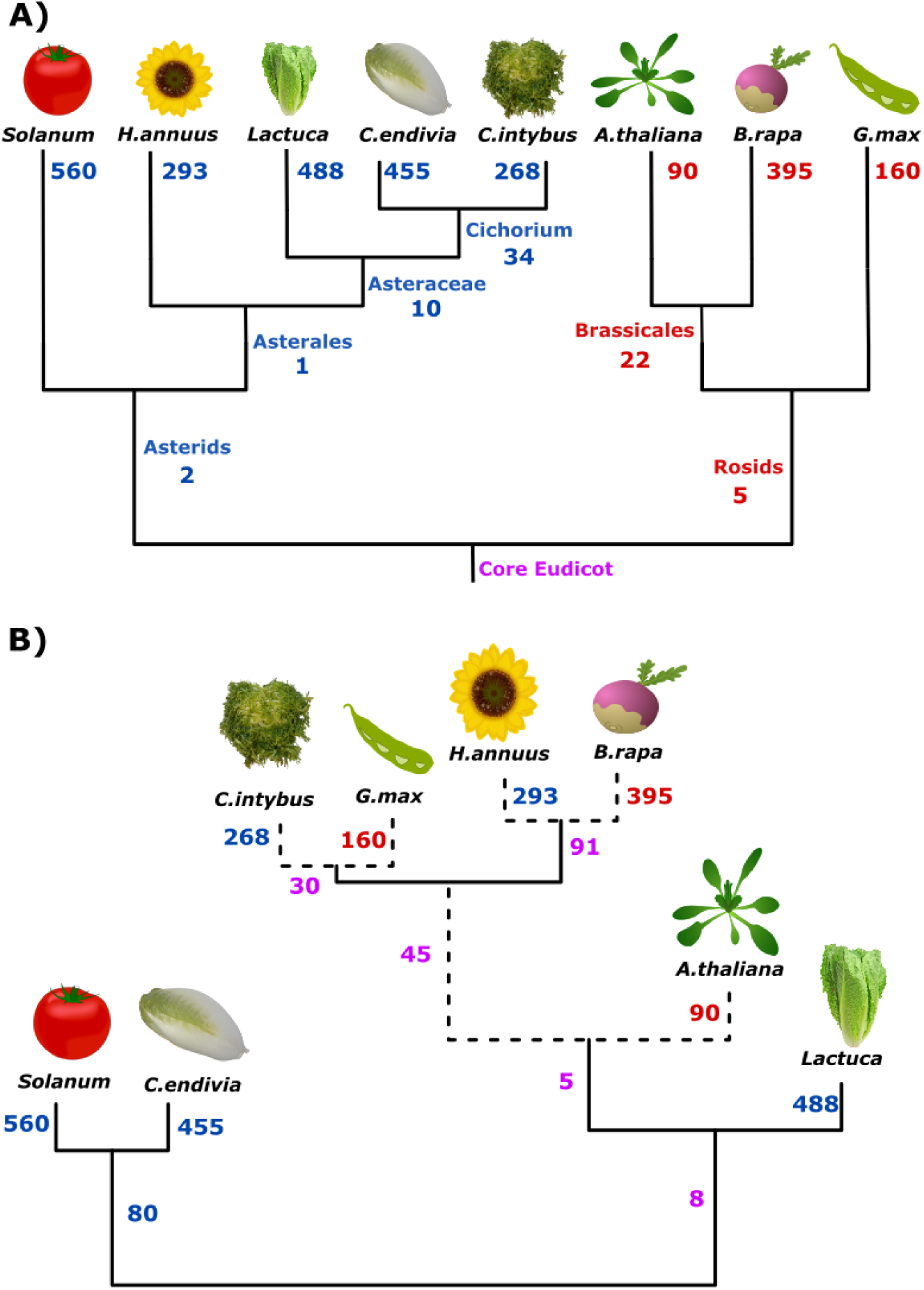
The Botrytis genes associated to virulence across eight eudicots do not track the hosts’ evolutionary history. The numbers of candidate genes associated to lesion area in red are for rosids, in blue for asterids, and in purple for the mixed association. A) Plant-driven evolution: Phylogenetic tree of the studied eudicot species with at each branch and node the number of candidate genes identified. B) Fungi-driven evolution: Hierarchical clustering of the candidate genes on the different hosts. Dashed lines represent non-significant branches.

As an alternative analysis, we used the candidate genes to test if the similarity in candidate virulence genes associated to the host species may show relationships that do not track the evolutionary phylogenetic relationships of the hosts. This test was done using a hierarchical clustering of the candidate genes effect across the different hosts (Figure 6B). Based on the fungal genetic variance, there is a significant discord with the plant phylogenetic relationship. Specifically, Botrytis virulence variation connects several Asterid-Rosid pairs including chicory (Asterids) clustering with soybean (Rosids) and sunflower (Asterids) clustering with turnip (Rosids) (Figure 6B). Similarly, tomato and endive identify a common set of 80 Botrytis candidate virulence genes (Figure 6B). These associations suggest that Botrytis may use similar virulence strategies across relatively unrelated host species. Such an observation might relate to the recognition of phytochemicals (Kuroyanagi *et al*., 2022) and activation of detoxification mechanisms that can be effective on phytochemicals from distantly related species (Bulasag *et al*., 2023; Wu *et al*., 2024). Plant defense phytochemicals often show convergent evolution in distantly related hosts like glucosinolates present in both the Brassicales and unrelated Drypetes (Pichersky & Lewinsohn, 2011; Negin & Jander, 2023). However, how Botrytis perceives the hosts and adapts its virulence strategies remains to be fully discovered.

While individual genes showed few common patterns across the diverse hosts (Figure 6), we proceeded to plot gene networks with potential virulence effects. We used the available Botrytis-Arabidopsis transcriptome dataset (Zhang *et al*., 2017) to plot the co-expressed candidate genes with correlation ρ>0.75. This approach revealed two main large networks (Figure 7). Network 1 is composed of a majority of genes annotated as integral membrane components, active in transmembrane transport or vesicular activities. Network 2 is composed of genes associated to the ribosome, translation, and protein folding activities. This analysis showed that the networks link candidate genes associated with diverse hosts. The shared virulence genes (identified in 4 or more hosts) are spread across the networks, linked by genes identified in one to three hosts (Figure 7). This suggests the potential for candidate genes identified in a few hosts to influence virulence by modulating the function of key *B. cinerea* networks.

**Figure 7:**
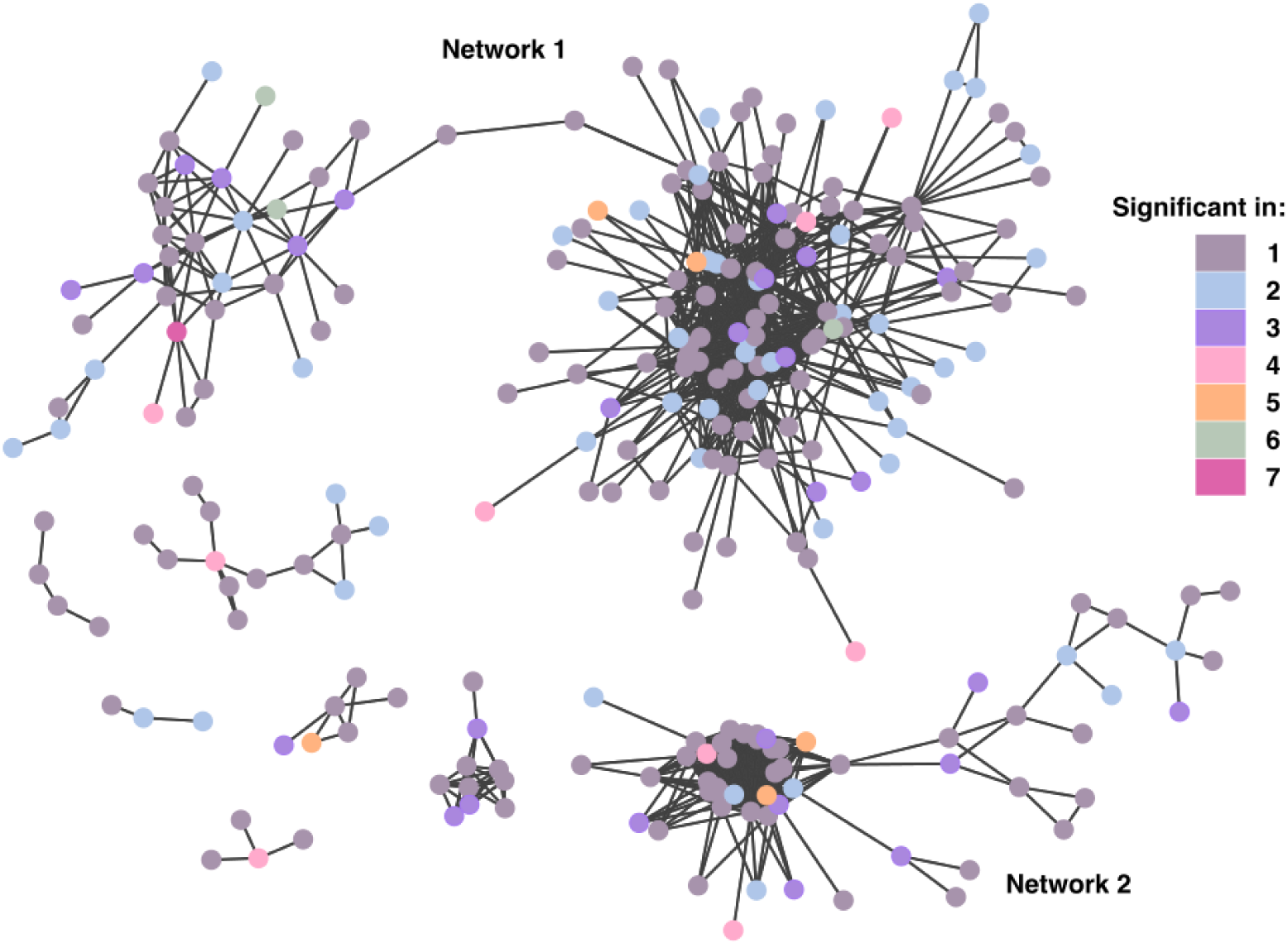
The Botrytis genes associated to virulence are active within co-expression networks. Candidate gene co-expression networks were calculated from *A.thaliana* transcriptome (Zhang *et al*., 2019). The vertices represent correlation ρ>0.75 at p<0.05. The dots represent candidate genes colored according to the number of hosts they were associated to.

## Discussion

In support of previous findings that *B.cinerea* genome was enriched in virulence factors in comparison to other fungal genomes (Amselem *et al*., 2011), the GWAS revealed that about one-third of the genome (Table S2) may influence Botrytis virulence and ability to infect different hosts. The candidate SNPs and genes found across all 16 core chromosomes had predominantly small effect sizes and were evenly distributed across the chromosomes. This distribution suggests a large influence of the core genome on the host range and does not support a bipartite architecture (Dong *et al*., 2015) with virulence-enriched genome areas (e.g. two-speed genome model) in this pathogen (Figures 2, S4). The genome-wide distribution of SNPs and genes associated to virulence is also consistent with the genome scans for evolutionary selection (Van Kan *et al*., 2017; Atwell *et al*., 2018; Mercier *et al*., 2021), which did not detect hotspots. Overall, this suggests that polymorphism spread across the core genome shapes the genetics of Botrytis virulence across the diversity of hosts tested.

In contrast to other systems, the candidate SNPs were not found on the accessory chromosomes. These accessory chromosomes contain only a few genes (less than 20) (Van Kan *et al*., 2017) with no significant SNPs in the coding sequences (Atwell *et al*., 2018). Further, the genes on the accessory chromosomes are expressed less frequently in planta than genes on the core chromosomes (Zhang *et al*., 2019). This differs from accessory chromosomes in other fungal species that contain host-specific virulence factors and toxin biosynthesis gene clusters (Bertazzoni *et al*., 2018) and have enriched expression in planta (Feurtey *et al*., 2020). While current data does not support a major role in the host range for these accessory chromosomes, further work with high-quality long-read genomes is needed to identify the full diversity of these chromosomes in the species. This is key in light of the model that the accessory chromosomes could be reservoirs of transposable elements contributing to small RNA production (Simon *et al*., 2022).

Interestingly, the GWAS identified mainly genes/SNPs influencing virulence in a small subset of hosts (Figures 5 and 6). Moreover, the identified genes are not unique to *Botrytis cinerea* but predominantly shared with other Botrytis species (Figures 4, S7), including species that specialize in infecting monocot plants (Valero-Jiménez *et al*., 2019; Valero-Jiménez *et al*., 2020). This suggests that virulence and host range variation in *Botrytis cinerea* may not primarily rely on creating new genes or mechanisms, but rather on adjusting existing ones. Differential gene expression in response to infection was shown across different hosts in *Sclerotinia sclerotiorum* (Allan *et al*., 2019; Kusch *et al*., 2021) and within Arabidopsis in Botrytis. In Botrytis, eQTL analyses further revealed that the transcription plasticity is controlled via distantly acting hotspots (trans-regulation) (Soltis *et al*., 2020; Krishnan *et al*., 2023) rather than local control (cis-regulation) like in *Zymoseptoria tritici* (Abraham & Croll, 2023). Heterochromatin accessibility (Zhang *et al*., 2021) and transposable elements (Simon *et al*., 2022) might also play a role in adapting virulence to various hosts.

When considering that the GWAS-identified genes are part of gene co-expression networks (Figure 7), it suggests a model where the genetic variation may impact virulence and host range through a set of gene networks facilitating the pathogen’s interaction with diverse hosts. The resulting host-Botrytis interactions are complex, as GWAS-identified genes/SNPs in these gene networks do not closely follow the hosts’ evolutionary relationships. Few genes align with the host phylogeny and may be adapting to evolutionary changes amongst the hosts. However, the majority of genes do not track the host phylogeny, with sets of genes shared among pairs of asterids/rosids hosts (Figure 5). This raises the possibility that the variation in these genes may not respond specifically to the hosts’ evolutionary changes but instead may function as a form of bet-hedging (de Jong *et al*., 2011; Haaland *et al*., 2020). This blend of Botrytis genes tracking host evolution and other genes potentially as bet-hedging could result from fluctuations in the host diversity. In locations with a long-term stable agricultural monocrop, Botrytis may have the opportunity to adapt genes/mechanisms to better infect specific hosts or plant families. This was documented in grapevine and tomatoes with the signature of positive selection on cell wall degradation enzymes and oxidative stress genes (Mercier *et al*., 2021). However, in environments with fluctuating host presence, genes/mechanisms that provide an advantage to diverse hosts might be favored.

While the GWAS did not identify genes affecting all eight hosts, it identified a small subset of genes affecting six or more hosts. Among those genes potentially acting as general virulence mechanisms (Figures 5, 6), 11 genes may be related to the fungal allorecognition and immune system (Table S2). These proteins composed of heterokaryon incompatibility, NB-ARC, NACHT, WD40, Tetratricopeptide repeat, and Ankyrin-repeats domains are associated to the regulation of cell death (Arshed *et al*., 2023; Daskalov, 2023).

To conclude, to successfully infect thousands of plants (Elad *et al*., 2016; Singh, R *et al*., 2023), *Botrytis cinerea* probably relies on a large portion of its genome involved in a multi-layer quantitative system. Some virulence mechanisms might be host-specific while others might contribute to Botrytis virulence across multiple hosts, potentially unrelated. This complex quantitative system is also highly redundant as shown by the resilience of phytotoxic machinery to serial knockouts (Leisen *et al*., 2022) and the four different strategies for the detoxification of a single phytoalexin (You *et al*., 2024). To precisely map how Botrytis recognizes the hosts, how the hosts impact the virulence strategies, and how the natural genetic diversity of Botrytis strains contributes to the virulence and host range will require a massive transcriptome sequencing effort and validation of gene networks rather than individual genes.

## Data availability

Correspondence and requests for materials should be addressed to. The datasets and R codes are available on Dryad: https://doi.org/10.5061/dryad.j6q573npm

To download the archive: https://datadryad.org/stash/share/UTNBJJp_D92KlsE51usJgyuxRynlP3qiw1RADV0G418

## Acknowledgments

This work was supported by the NSF award IOS 2020754 and USDA award 2019-05709 to DJK.

## Authors contribution

Data analysis: C.C.; Funding acquisition: D.J.K., Investigation: C.C., D.J.K.; Writing: C.C., D.J.K.

---

**Figure S1:**
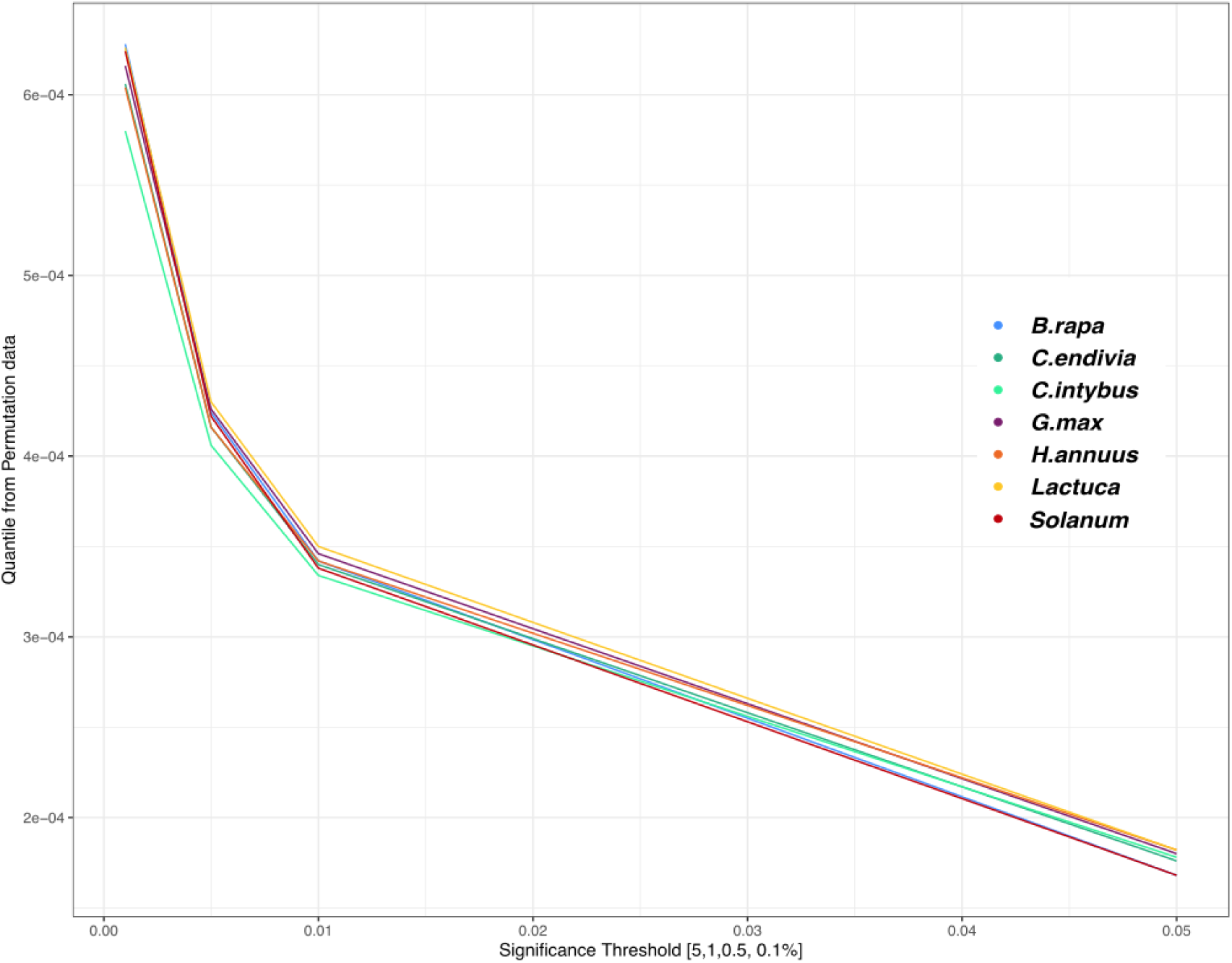
Permutation-based significance thresholds for the posterior inclusion probability (PIP). For the lesions on each species (phenotype), the significance thresholds were estimated by running the BSLMM model with ten random permutations of SNPs positions. PIP values larger than 1.7 x10^-4^ are equivalent to 5% chance of a false positive, while 3.4 x10^-4^ and 4.2 x10^-4^ are respectively at 1% and 0.5% thresholds.

**Figure S2:**
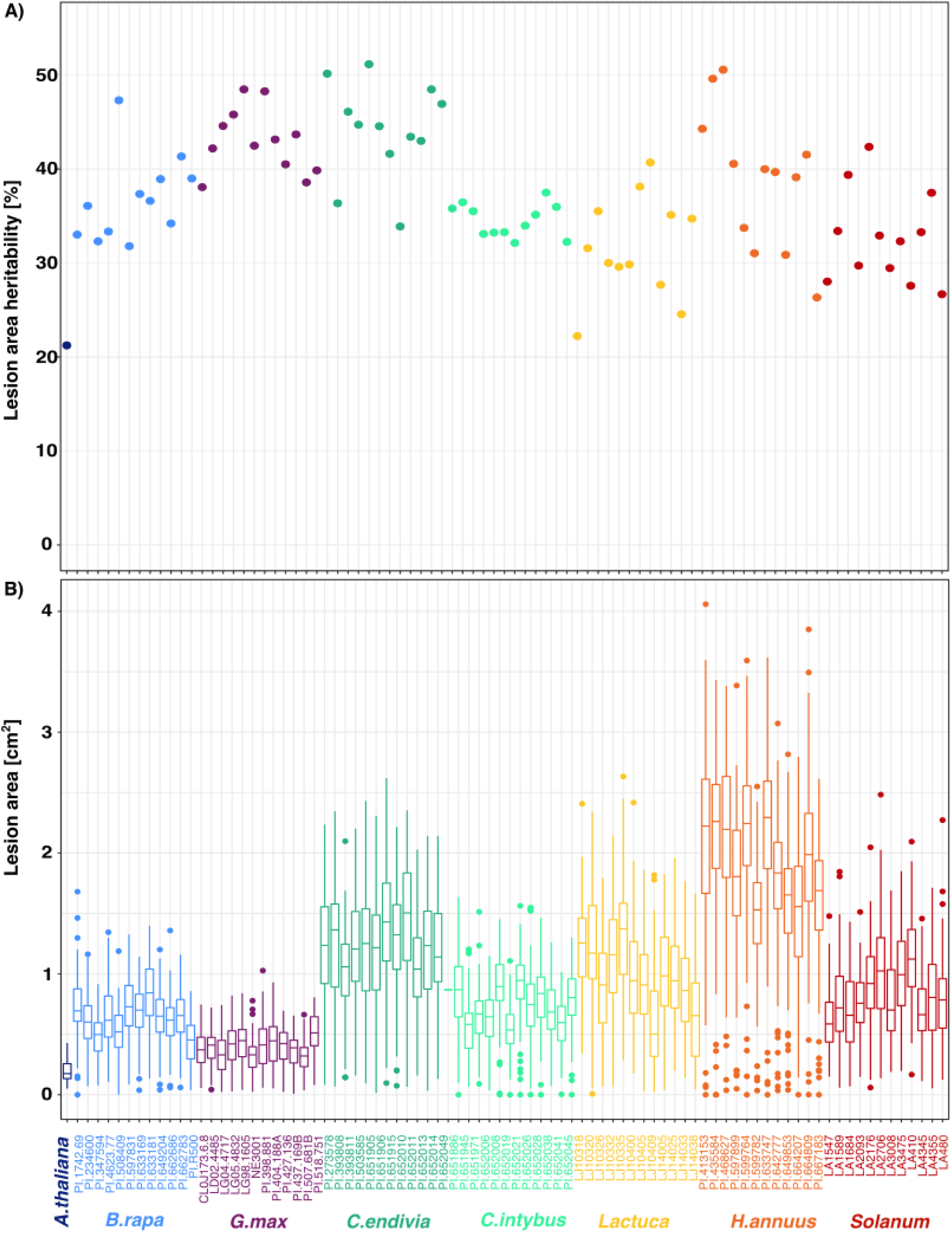
Overview of the phenotypic data (Caseys *et al*., 2021) used to conduct the GWAS. A) Genetic heritability of the lesion area of 96 isolates of Botrytis on 85 plant genotypes colored by host species. The heritability was calculated as the percentage of variance in lesion area explained by Botrytis isolates. B) Variation in lesion area at 72 hours post inoculation across the 96 Botrytis isolates for each plant genotype.

**Figure S3:**
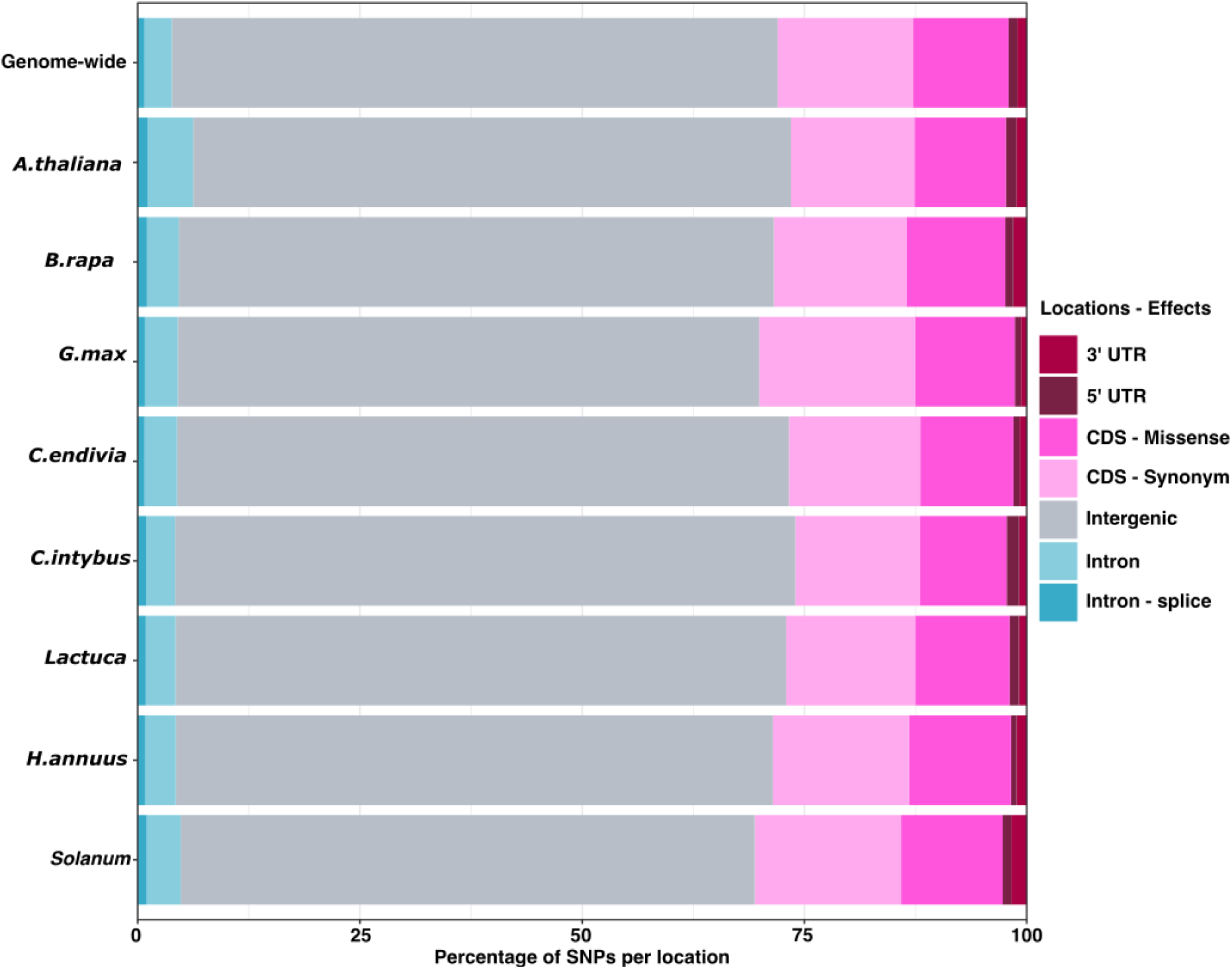
Significantly associated SNPs (LFSR<0.05) show no enrichment for genomic features. Count SNPs for each host species categorized based on their genomic locations and effects. No significant deviation from the genome-wide distribution was detected.

**Figure S4:**
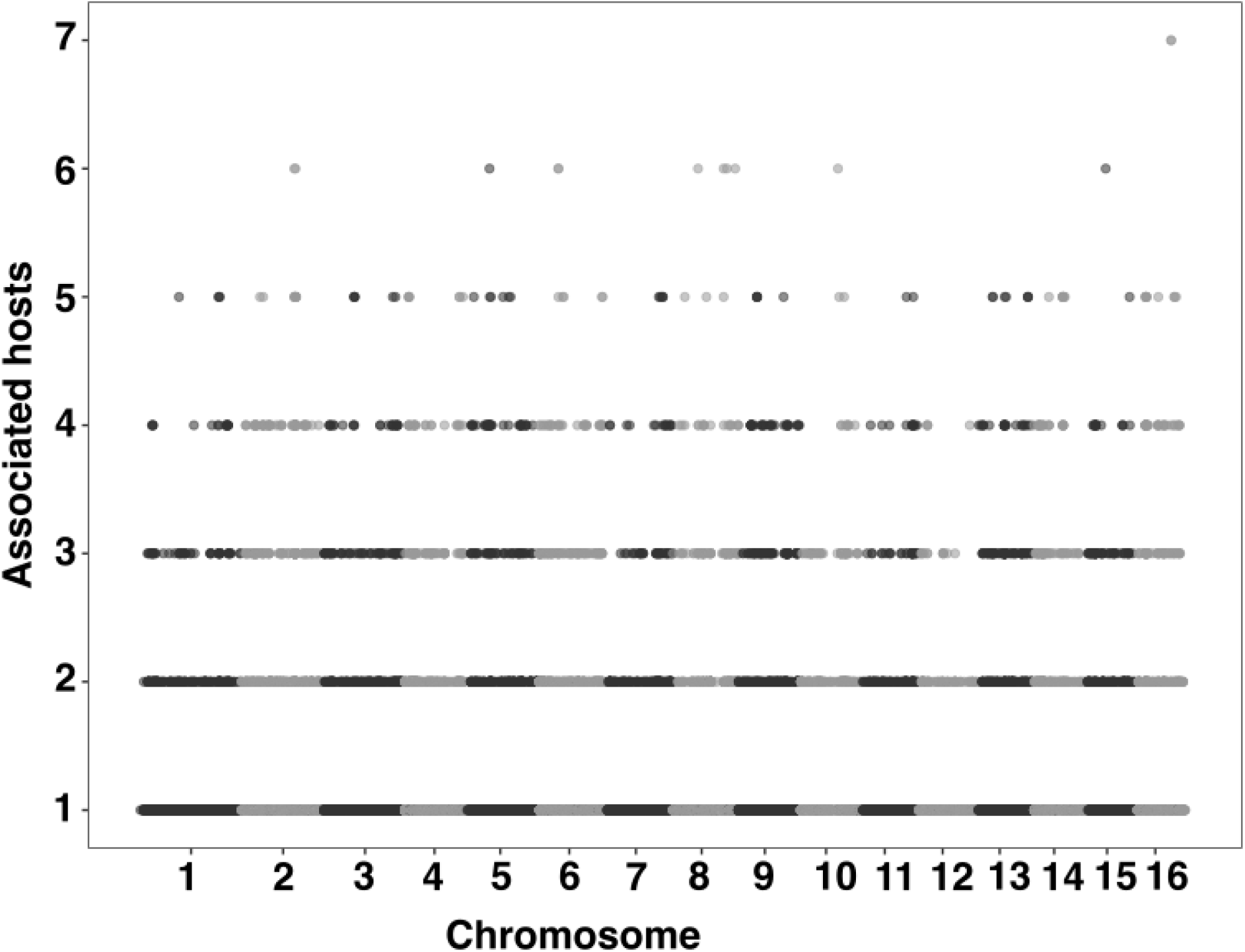
Genome-wide distribution of SNPs significantly (LFSR<0.05) associated to one to seven hosts. The 16 chromosomes of *B. cinerea* are delimited by shades of grey.

**Figure S5:**
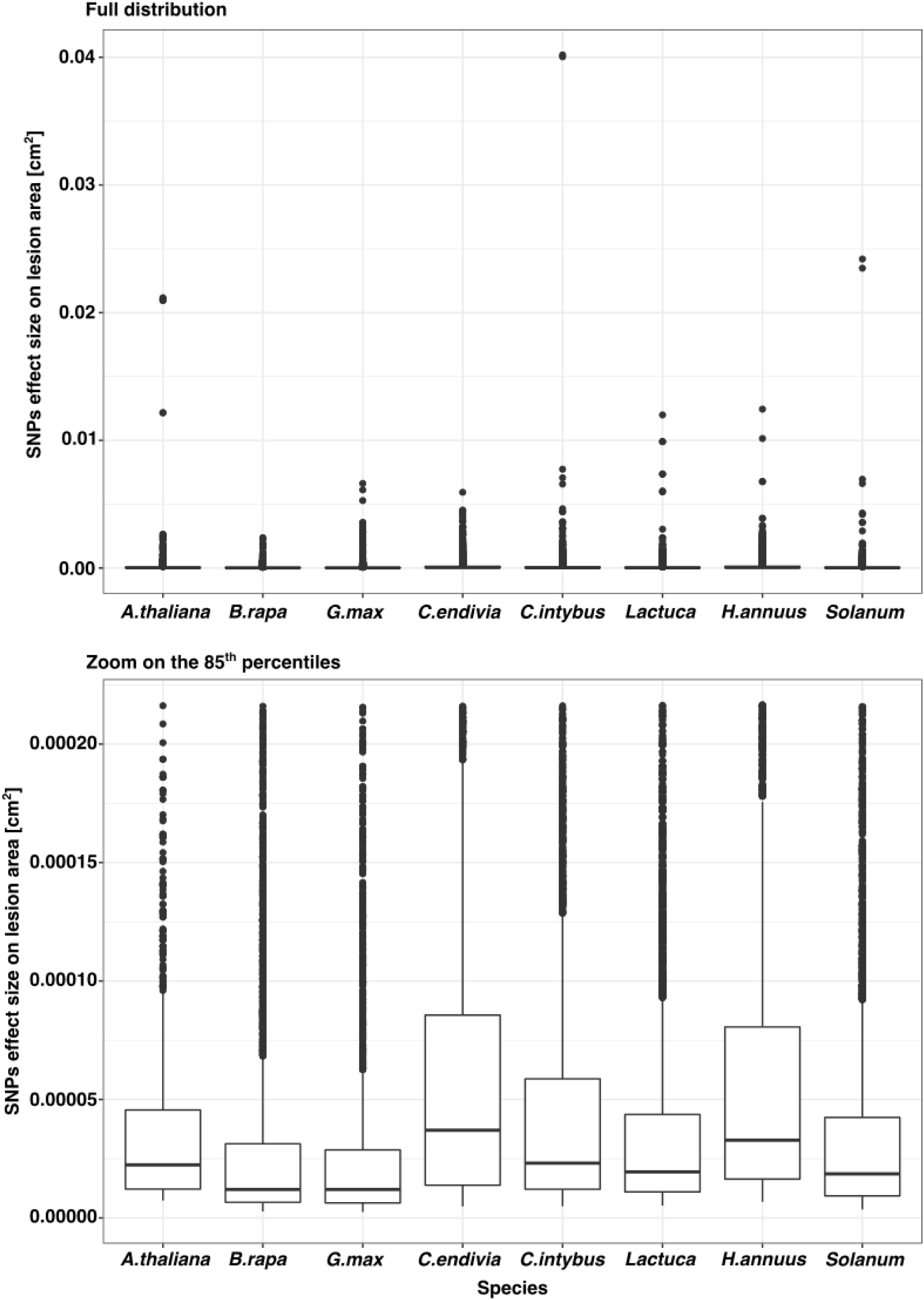
Distribution of the effect size of significantly associated (LFSR<0.05) SNPs to lesion area on eight host species. Given the large range of effect sizes, both the full distribution and zoom to the 85^th^ percentile are provided.

**Figure S6:**
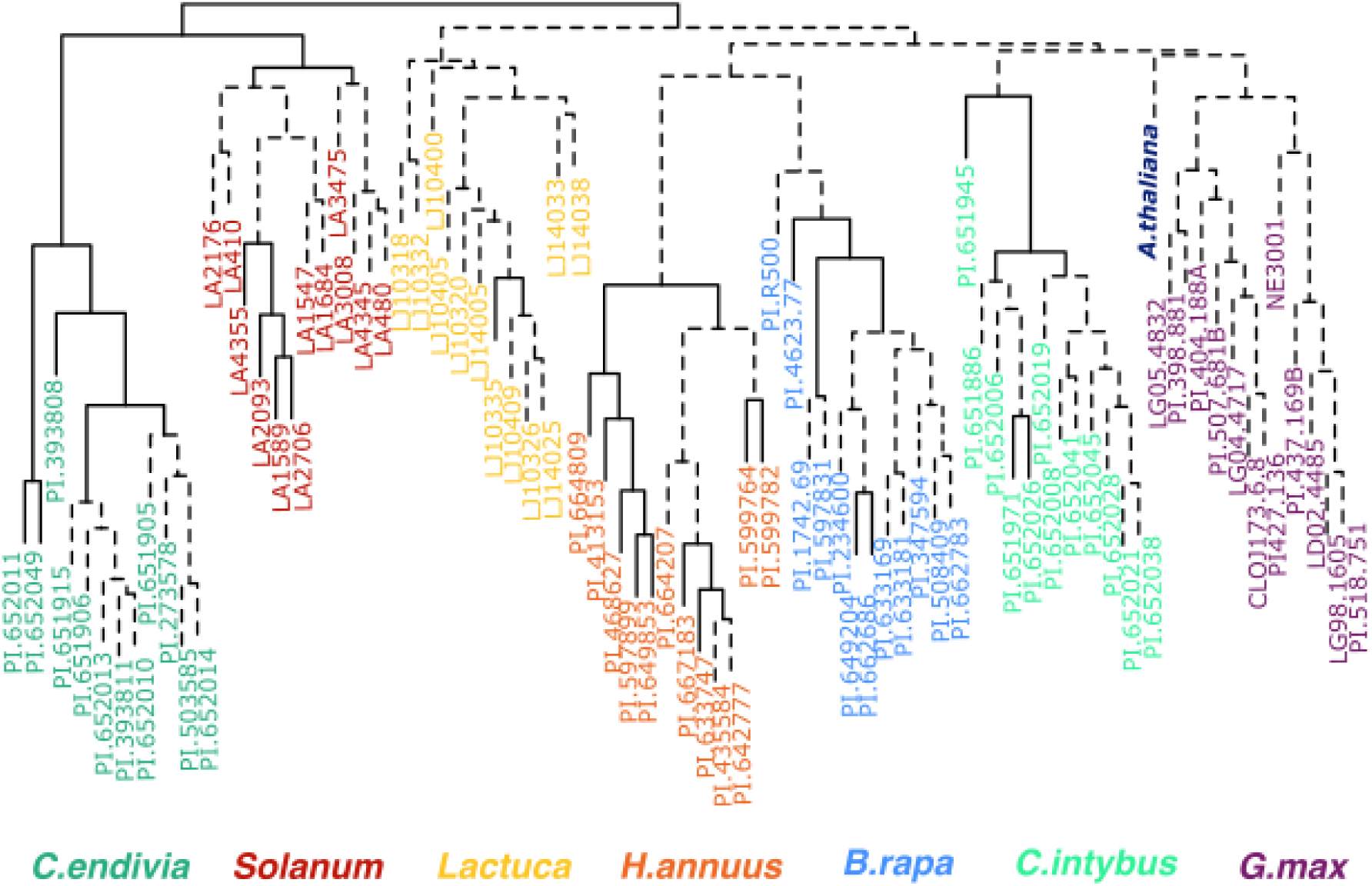
Hierarchical clustering of significantly associated SNPs (LFSR<0.05) across 85 host genotypes. Branch length represents the correlation distances between genotypes based on the significance status of each SNP on each genotype. Dashed lines represent non-significant (p-val>0.05) branches from bootstrap resampling. The genotype names are colored based on the host species.

**Figure S7:**
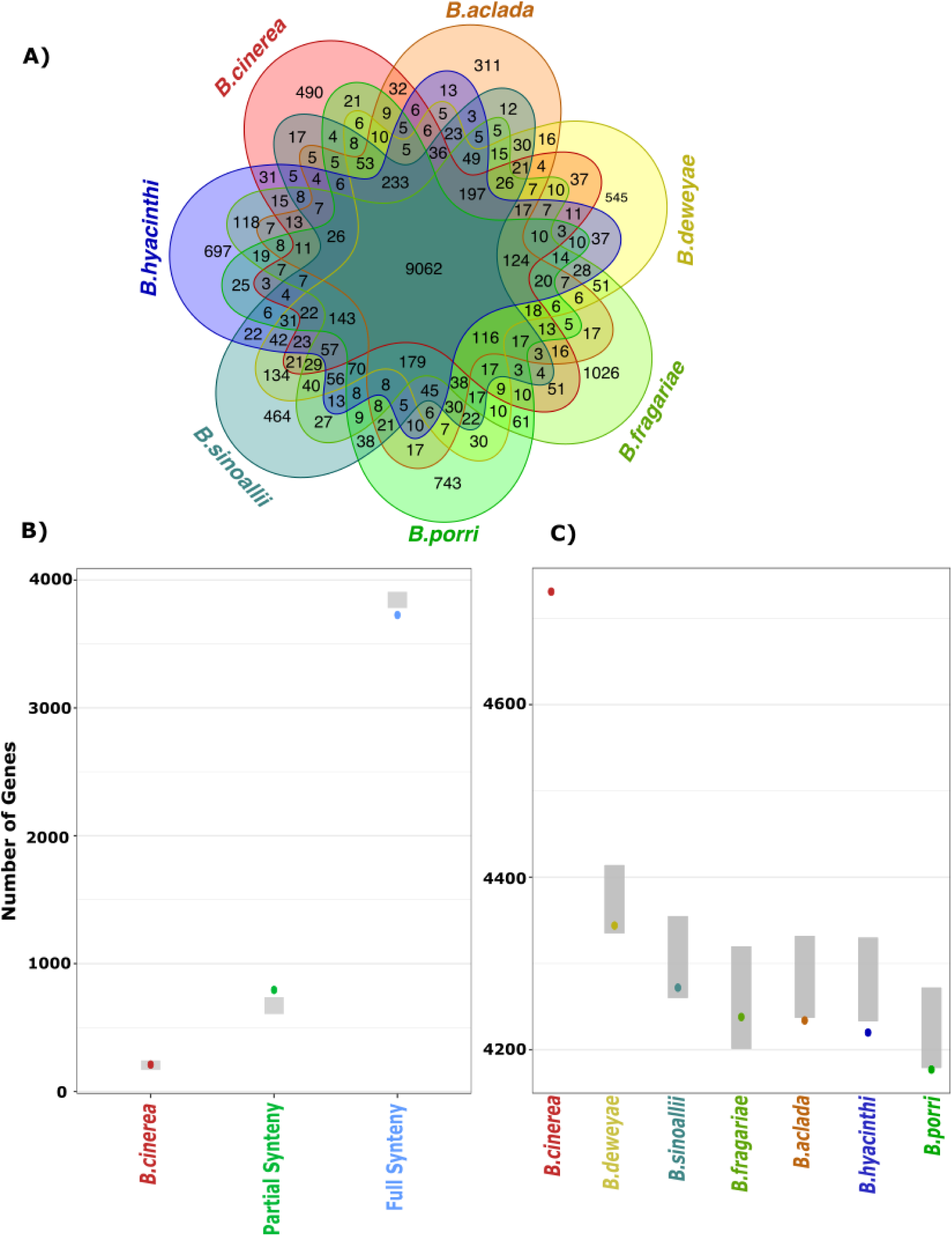
Synteny among seven Botrytis species. A) Venn diagram with the count of orthologs between *B.cinerea* (B05.10 reference genome) and six other Botrytis species. B) Number of candidate genes identified by GWAS (dots) with no (identified only in *B.cinerea*), partial (identified in 2 to Botrytis 6 species), or full synteny (identified in 7 Botrytis species). The grey zones represent the minimum and maximum range observed in the thousand random gene sets. C) Number of candidate genes identified by GWAS (dots) with orthologs in the Botrytis species. The grey zones represent the minimum and maximum range observed in the thousand random gene sets.

